# Elucidating the Bell-Shaped Dependence of Protein Translation Activity on EF-Tu Concentration in a Reconstituted Cell-Free System Using a Mechanistic Model

**DOI:** 10.64898/2026.04.17.719328

**Authors:** Shunnosuke Ban, Yusuke Himeoka, Ako Kagawa, Yoshihiro Shimizu, Tomoaki Matsuura, Chikara Furusawa

## Abstract

Protein synthesis in cell-free protein synthesis systems often exhibits non-intuitive input-output relationships. In the PURE system, a reconstituted cell-free system, protein production peaked at low elongation factor Tu (EF-Tu) concentrations and decreased at higher concentrations, resulting in a characteristic bell-shaped profile. Here, we investigated the origin of this behavior using a detailed mechanistic model of translation in the PURE system, designated as ePURE, which describes reaction dynamics of hundreds of molecular species and reactions. Our computational analysis suggested that excess EF-Tu sequesters the initiator tRNA (tRNA^fMet^) into non-productive EF-Tu·GTP·Met-tRNA^fMet^ complexes, thereby depleting the pool of initiator tRNA available for translation initiation. This suppression arises from competition for a limited molecular resource rather than from direct inhibition. Based on this mechanism, we predicted that increasing the concentrations of tRNA^fMet^ and methionyl-tRNA formyl-transferase would eliminate the bell-shaped dependence, and experimentally confirmed this prediction. Under these modified conditions, the bell-shaped response disappeared and protein production was enhanced. These findings demonstrate how mechanistic computational models can reveal hidden constraints underlying non-intuitive input-output relationships in complex biochemical networks and guide the rational optimization of cell-free protein synthesis systems.

## 1 Introduction

Cells carry out essential processes such as transcription, translation, and metabolism through intricate networks of chemical reactions (*1*). These reactions are precisely regulated and flexibly adjusted in response to environmental cues, including nutrient availability, temperature, and stress (*2* –*4*). This regulation is inherently difficult because many molecules participate simultaneously in multiple reactions, creating competition and interdependence among pathways (*5*). Consequently, increasing the supply of a substrate or cofactor does not necessarily enhance the production of a desired molecule if the same input is required by competing reactions, or if downstream products inhibit the target pathway (*6*, *7*). These non-intuitive input-output relationships highlight the complexity of biochemical networks, in which shared substrates and products can lead to unexpected outcomes. Understanding these dynamics is crucial for elucidating the robustness and adaptability of living systems and for advancing synthetic biology approaches that reconstruct and engineer complex cellular functions *in vitro*.

Experimental analysis of input-output relationships in living cells remains challenging. *In vivo* systems contain numerous interacting components, including unidentified ones, which obscure the specific reactions or parameters that govern the behaviors of interest. To address this limitation, we employ an *Escherichia coli*-based reconstituted cell-free translation system, known as the PURE system (*8*). This system consists of a set of purified factors that are sufficient for efficient protein synthesis. Experimentally, the PURE system enables quantitative exploration of input-output relationships by varying the initial concentrations of components (inputs) and measuring peptide production (outputs). Previous studies that systematically altered the initial concentrations of PURE components reported several non-intuitive production profiles, including bell-shaped dependencies in which peptide production increases with a factor’s concentration up to a certain threshold and then decreases (*9*). Such bell-shaped dependencies are characteristic signatures of competition and inhibition within complex reaction networks.

Unraveling the detailed mechanisms underlying these input-output relationships requires extensive experiments under diverse conditions, along with time-resolved measurements of both final products and intermediate species. However, experimentally manipulating reaction parameters and monitoring time-series data for all components is extremely demanding. In such cases, computational modeling offers a powerful alternative (*10* –*14*). For the PURE system, a detailed mechanistic model of the PURE system, which we refer to here as ePURE, has been developed (*15*). It comprises hundreds of components and reactions and simulates the translation process using ordinary differential equations. This model allows the time evolution of all components in the PURE system to be calculated, making it possible to systematically investigate how variations in initial concentrations influence translation outcomes (*15* –*17*).

In this study, we aim to elucidate the mechanisms behind complex input-output relationships in translation systems, focusing on the bell-shaped dependence of peptide production on the input concentration of elongation factor Tu (EF-Tu). EF-Tu is essential for the peptide elongation step, yet our PURE system experiments revealed a pronounced bell-shaped dependence of protein production on EF-Tu concentration that lacked a clear mechanistic explanation. Because the original ePURE model could not reproduce this phenomenon quantitatively, we first refined the kinetic parameters by fitting the model output to experimental data. Analysis of the adjusted parameters allowed us to identify the reaction network responsible for the bell-shaped dependence and reveal a strategy for eliminating it. Guided by this finding, we designed an experimental setup to modulate this dependence and successfully eliminated the bell-shaped response. Additionally, the peptide production further increased owing to the elimination of bell-shaped dependence. These results demonstrate that mechanistic computational models can not only explain counterintuitive input-output relationships in complex biochemical networks but also guide rational redesign of cell-free systems to achieve desired performance.

## 2 Results and Discussion

### 2.1 Non-monotonic EF-Tu Dependence of Peptide Synthesis in the PURE System

We quantified how the amount of peptide production depends on the initial EF-Tu concentration in the PURE system. A reaction mixture containing 67 defined components was reconstituted, and only the initial EF-Tu concentration was varied across five levels (0, 5, 25, 50, and 65 µM), with all other components held constant (see Methods and Table S1). Green fluorescent protein (GFP) (sequence given in Table S2) was used as a reporter, and its fluorescence increase was recorded over time. To compare the experimental data with the ePURE model, which explicitly describes the synthesis of the Met-Gly-Gly tripeptide, we converted the GFP fluorescence intensity into the corresponding amount of synthesized peptide (details in Methods).

As expected, no peptide was produced in the absence of EF-Tu, which is essential for the transfer of aminoacyl-tRNAs to the ribosomal A site. However, when EF-Tu was supplied at concentrations above 5 µM, the peptide production decreased as EF-Tu concentration increased (Figure 1A). To clarify this trend, the amount of synthesized peptide at the endpoint (1,560 s) was plotted against the initial EF-Tu concentration (Figure 1B). Although increasing EF-Tu would normally be expected to enhance translation until another component becomes rate-limiting, our experiments showed the opposite: higher EF-Tu concentrations reduced the amount of peptide production. Rather than approaching saturation, the EF-Tu dependence of the peptide synthesis exhibited a bell-shaped profile. This behavior provides a counterintuitive example of a non-monotonic input-output relationship in a biochemical network and forms the central focus of the present study.

**Figure 1:**
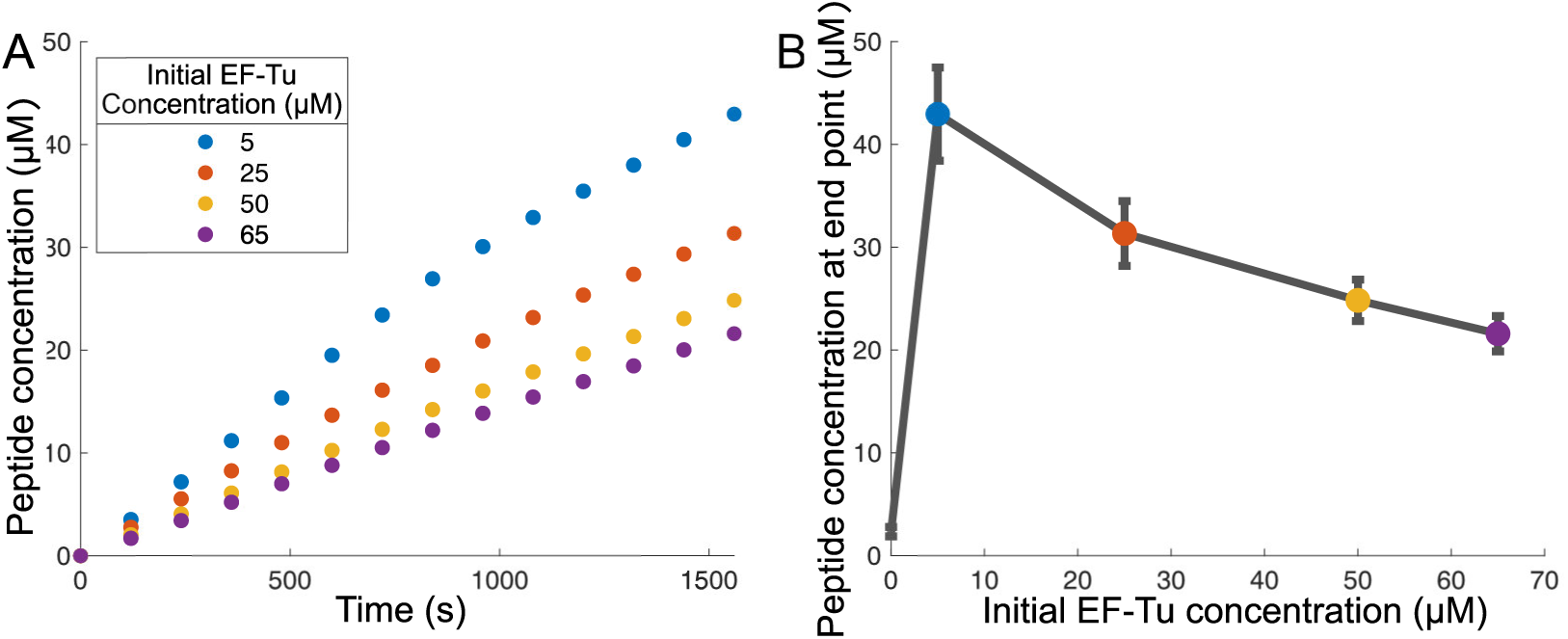
Experimental results of the effect of initial EF-Tu concentration on peptide synthesis. (A) Time course data for each initial EF-Tu concentration. (B) Peptide concentrations at the endpoint (1,560 s). Error bars represent the standard deviation (s.d.) of four replicate experiments (n=4).

### 2.2 Simulations with Kinetic Parameters fitted by a Genetic Algorithm

To uncover the mechanism underlying the bell-shaped dependence observed in Figure 1, we sought to reproduce the experimental results by numerically simulating the PURE system. The ePURE model describes translation in the PURE system with ordinary differential equations and includes 205 variables for the concentrations of compounds and their complexes and 481 reactions among them. Using literature-derived kinetic parameters, the model calculates the time evolution of all concentrations.

Original ePURE parameters gave rise to a protein synthesis time course that reaches a steady-state in a few minutes, consistent with experimental data. However, with the original ePURE parameters, the simulated peptide production was substantially slower than observed experimentally (Figure 2B). Before analyzing the bell-shaped dependence, we therefore addressed this quantitative gap between the simulation and the experiment. We reasoned that the discrepancy likely arose because the default kinetic parameters, previously collected from the literature, were measured in highly purified forms and under conditions different from the PURE system, and may not necessarily capture the complexity of the PURE system.

**Figure 2:**
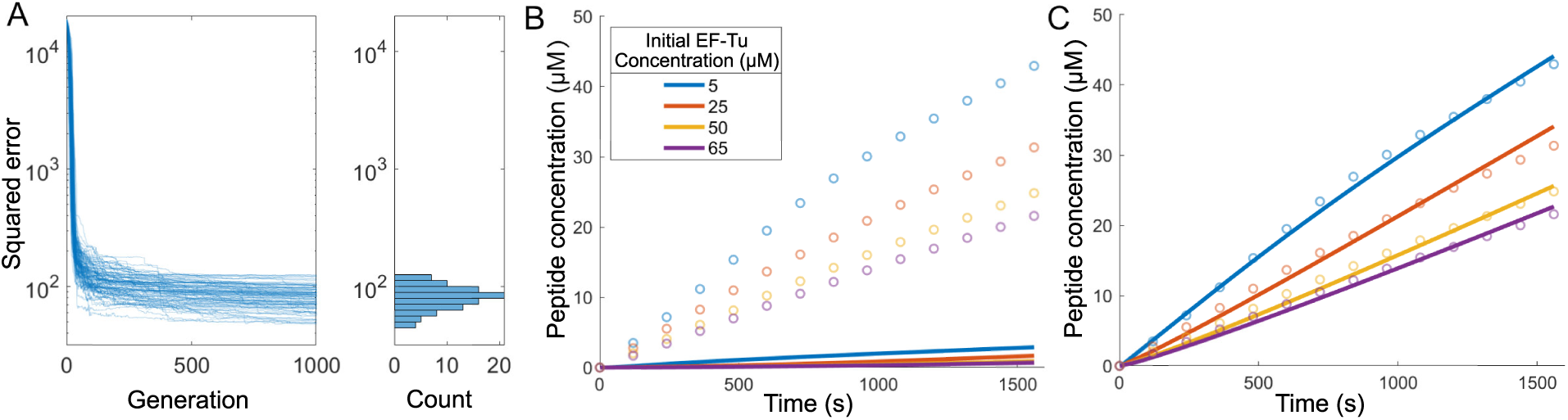
Parameter fitting of ePURE by GA. (A) Change in the squared error between experimental data and simulations over GA generations (left) and its distribution in the final generation (right). GA performed 100 runs of 1,000 generations each. Changes in penalties and values of the objective function are provided in Figure S1. (B) Simulation results using the original kinetic parameters and the initial component concentrations of the ePURE system. The dots and solid lines represent the experimental observations and the simulated results, respectively. The initial concentrations used in the simulation corresponding to the experiment are shown in Table S3. (C) A representative simulation result obtained using fitted kinetic parameters.

To reproduce the experimental behavior, we then attempted to modify the ePURE’s kinetic parameters. Previous studies indicated that varying individual parameters, even over wide ranges, had a limited impact on translation rate, suggesting that single-parameter tuning is insufficient (*16*). Reproducing PURE dynamics would instead require coordinated adjustments to multiple parameters. We employed a genetic algorithm (GA) to estimate parameters so as to reproduce the experimental time courses, because GAs efficiently explore high-dimensional parameter spaces and are less prone to local optima than gradient-based methods in nonlinear dynamical systems with many variables and parameters. The objective function minimized the sum of squared errors between simulated and measured peptide concentrations (see Methods for details). To discourage unrealistic parameter drift, we added a penalty that increases with the magnitude of deviation from the default values.

Figure 2A shows the evolution of the objective function across 100 independent runs, including the generation-wise decrease in squared error (penalty excluded) and the distribution of errors in the final generation. GA fitting markedly reduced the mismatch relative to the original parameter set. In later generations, errors plateaued, consistent with convergence to local optima. The final error distribution also indicates substantial diversity among the best-performing solutions.

To assess the fidelity of simulations using the fitted parameters, we examined a representative run selected from the GA-derived sets (Figure 2C). Simulations with the original parameters deviated strongly from the data, most notably by underestimating the amount of peptide production (Figure 2B). In contrast, the GA-fitted parameters increased the peptide production while retaining the bell-shaped dependence on the initial EF-Tu concentration. We found that this non-monotonic dependence was consistently reproduced across all GA-fitted simulations.

### 2.3 Mechanism of Bell-Shaped Behavior: Excess EF-Tu Depletes tRNA^fMet^

The PURE system comprises a complex reaction network, which makes the mechanism of the bell-shaped dependence difficult to dissect. Figure 3A presents a schematic of the translation subnetwork involving EF-Tu, highlighting the relevant reactions in the initiation and elongation steps. Initiation and elongation require the corresponding aminoacyl-tRNA (aa-tRNA). Initiation requires formylated methionine. After formylation by methionyl-tRNA formyl-transferase (MTF) using 10-formyltetrahydrofolate (FD), fMet-tRNA^fMet^ binds to IF2·GTP to form the IF2·GTP·fMet-tRNA^fMet^ complex, which is used for initiation (*18* –*21*). For elongation, EF-Tu is a key elongation factor that delivers aa-tRNAs to the ribosome (*22* – *25*).

**Figure 3:**
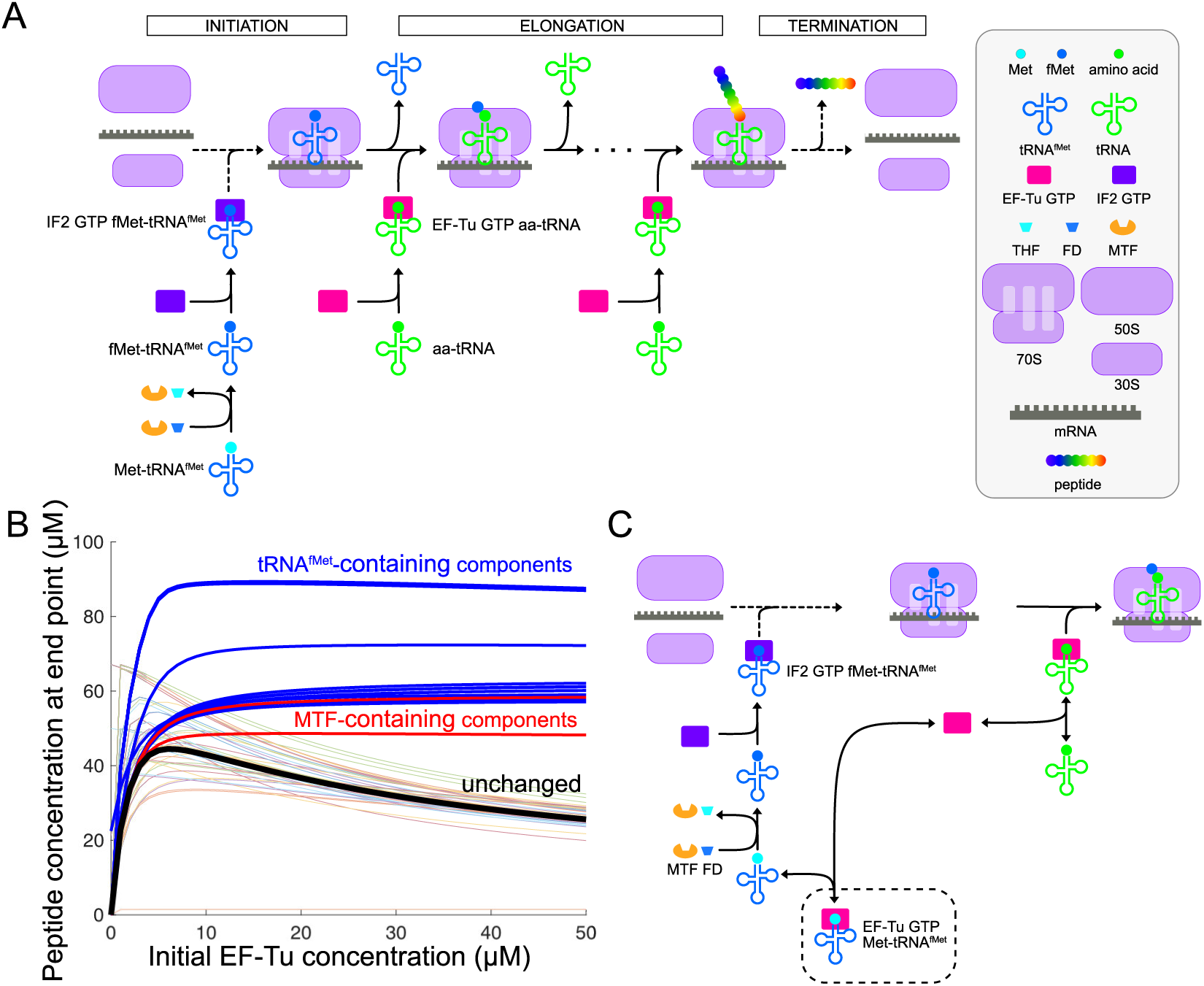
Mechanism underlying the bell-shaped dependence of the peptide production on EF-Tu concentration. (A) Schematic illustration of reactions in protein translation. For initiation, the methionine corresponding to the start codon is formylated, a reaction catalyzed by MTF, in which Met-tRNA^fMet^ receives a formyl group from 10-formyltetrahydrofolate (FD), a tetrahydrofolate derivative (THF). Subsequently, it binds to IF2·GTP to form the IF2·GTP·fMet-tRNA^fMet^ complex, which is used for initiation. For elongation, EF-Tu plays a central role by delivering aa-tRNA to the ribosome. (B) Numerical analysis of how increasing the initial concentration of each component affects the EF-Tu dependence of the peptide production. Each line shows the EF-Tu dependence obtained when the initial concentration of a single component was increased by 1 µM. The thick black line represents the result using the original initial concentrations. The thick blue and red lines show all the results obtained when the initial concentrations of the tRNA^fMet^-containing components and MTF-containing components were increased, respectively. Kinetic parameters were taken from a representative parameter set obtained by GA fitting. This trend is not limited to the 1 µM increment shown here; consistent results were obtained across other concentration increments (see Figure S2 for results with increments ranging from 0.05 µM to 5 µM). (C) Schematic illustration of EF-Tu-related reactions in protein translation. When EF-Tu·GTP binds Met-tRNA^fMet^, the resulting EF-Tu·GTP·Met-tRNA^fMet^ complex (indicated by dashed square) can no longer participate in elongation or initiation.

We hypothesized that increasing the initial EF-Tu concentration depletes specific components, and that exogenous supplementation of the depleted components would modulate the EF-Tu dependence of the peptide production. To test this, we performed simulations in which the initial concentration of each component was increased by 1 µM individually, allowing us to identify which components influence the bell-shaped response.

Figure 3B shows how the EF-Tu dependence of the peptide production at 1,560 seconds changed upon increasing the initial concentration of each component. Relative to the original bell-shaped curve shown by the thick black line, many components produced noticeable deviations, although many of the resulting curves remained bell-shaped. Among all components, we focused on cases in which the bell shape disappeared (e.g., blue or red curves). Components whose elevation eliminated the bell-shaped dependence clustered within a specific region of the reaction network; when we increased the initial concentration of complexes that contain tRNA^fMet^, the EF-Tu dependence changed markedly (thick blue lines). Increasing complexes that include MTF also abolished the bell shape (thick red line). These results indicate that initiation reactions involving tRNA^fMet^, shown in Figure 3C, are crucial for the observed dependence. Similar results were also obtained with other concentration increases (Figure S2).

Figure 3C presents a schematic of the EF-Tu-related part of the initiation and elongation steps. EF-Tu delivers aa-tRNA to the ribosome during the elongation step, but EF-Tu can also bind Met-tRNA^fMet^, the initiator tRNA (*26*). The EF-Tu·GTP·Met-tRNA^fMet^ complex is functionally inert in both initiation and elongation. Because no tRNA synthesis occurs in the PURE system, the total concentration of tRNA^fMet^ does not change. As EF-Tu increases, more Met-tRNA^fMet^ becomes sequestered in this non-productive complex, reducing the Met-tRNA^fMet^ available for formylation and initiation.

A bell-shaped dependence often implies that increasing an input factor initially promotes the reaction but subsequently suppresses it. In this system, the suppression does not arise from direct inhibition by EF-Tu. Instead, it emerges from competition for a limited resource: tRNA^fMet^. At high EF-Tu concentrations, the majority of Met-tRNA^fMet^ is trapped in EF-Tu·GTP·Met-tRNA^fMet^, leaving insufficient fMet-tRNA^fMet^ for initiation. Thus, the decline in peptide synthesis reflects competition between elongation-related binding of EF-Tu and initiation-specific processing of the initiator tRNA, rather than inhibitory activity of EF-Tu itself.

To evaluate this shortage more directly, we examined the simulated concentrations of complexes involving tRNA^fMet^. Because tRNA is not synthesized *de novo* in this system, the total amount of tRNA^fMet^ across free and bound states remained constant at 0.194 µM. Because these transient complexes influence the peptide synthesis rate at any given moment, their endpoint concentrations do not reflect their overall effect. Therefore, we evaluated their time-averaged concentrations over the entire simulation period (0–1,560 s) across different EF-Tu initial concentrations (Figure 4). As the initial EF-Tu concentration increased, EF-Tu·GTP·Met-tRNA^fMet^ (yellow region in Figure 4) came to dominate the total pool of tRNA^fMet^-related complexes, whereas IF2·GTP·fMet-tRNA^fMet^ (blue region) decreased to nearly zero. Because IF2·GTP·fMet-tRNA^fMet^ is the immediate precursor that engages the ribosome during translation initiation, its near depletion indicates that initiation becomes rate-limiting under high EF-Tu conditions (see Figure S3). Importantly, this mechanism was not specific to the representative fit shown here. Across all GA-fitted trials, excess EF-Tu consistently promoted the formation of EF-Tu·GTP·Met-tRNA^fMet^ complexes (Figure S4), depleted free initiator tRNA, and reduced initiation at higher EF-Tu concentrations.

**Figure 4:**
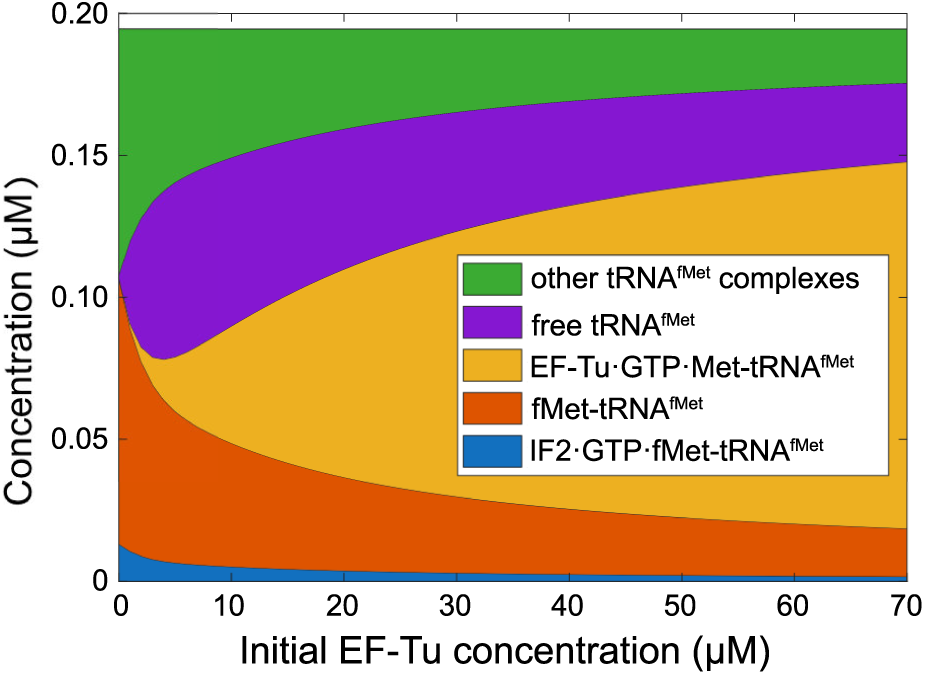
Time-averaged concentrations of tRNA^fMet^-containing complexes as a function of initial EF-Tu concentration. The stacked area plot illustrates the shifting distribution of these complexes. As the initial EF-Tu concentration increases, EF-Tu·GTP·Met-tRNA^fMet^ (yellow) becomes dominant, whereas IF2·GTP·fMet-tRNA^fMet^ (blue), which is required for translation initiation, decreases.

We further tested whether increasing the initial tRNA^fMet^ concentration would suppress the bell-shaped dependence. Using the same kinetic parameters as in Figure 2C, we increased only the initial tRNA^fMet^ to 1.94 µM. Upon increasing tRNA^fMet^, the decrease in the peptide production at higher EF-Tu concentrations disappeared, and the production remained comparable to that at 5 µM EF-Tu (Figure 5A). Consistently, the EF-Tu·GTP·Met-tRNA^fMet^ complex no longer dominated the total tRNA^fMet^ pool, and the concentration of the IF2·GTP·fMet-tRNA^fMet^ complex was stably maintained across the tested range of EF-Tu concentrations (Figure 5B). These findings indicate that, even under excess EF-Tu, a sufficient amount of fMet-tRNA^fMet^ can be maintained to support initiation, thereby preventing translation inhibition. Taken together, the ePURE simulations corroborate the proposed mechanism.

**Figure 5:**
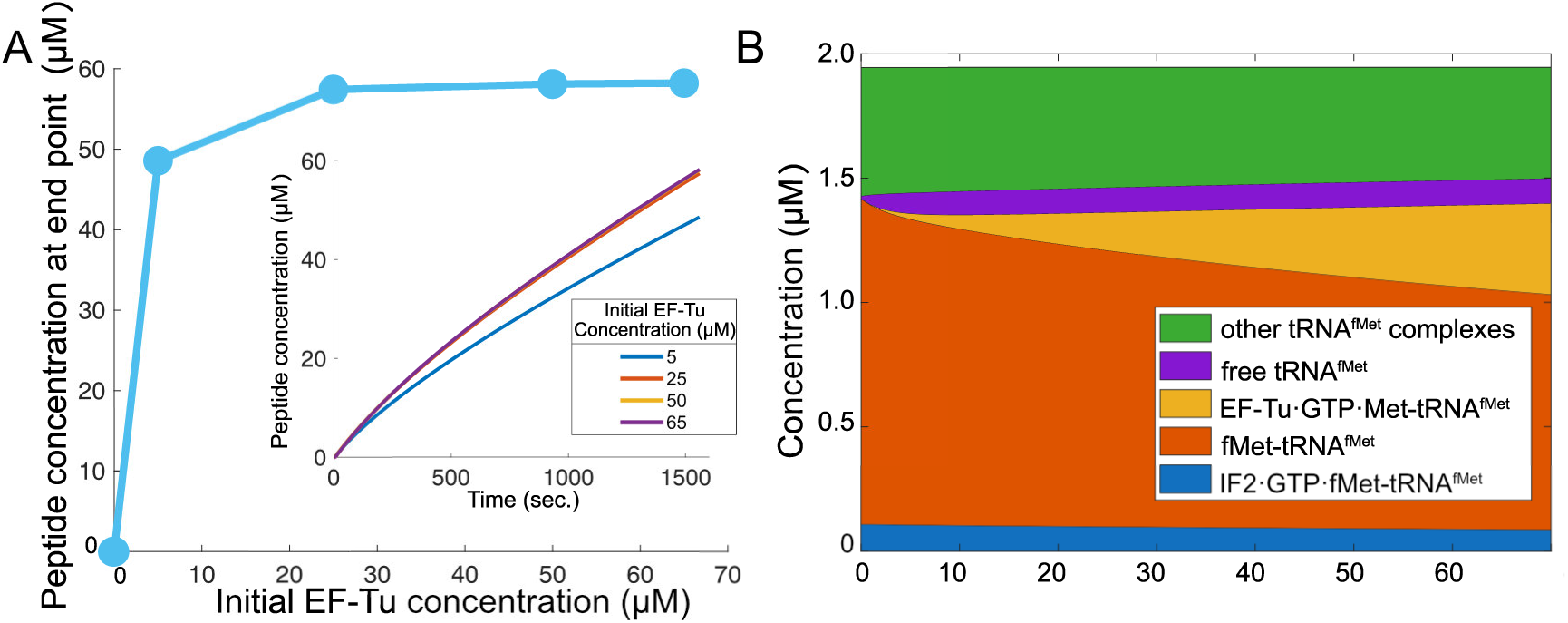
Simulation results with an increased initial tRNA^fMet^ concentration. (A) Peptide concentrations at endpoint (*t* = 1, 560*s*) with increased tRNA^fMet^ concentration to 1.94 µM (from the original 0.194 µM). The inset shows the time course of peptide concentration under these conditions. (B) Time-averaged concentrations of tRNA^fMet^-containing complexes as a function of initial EF-Tu concentration, with tRNA^fMet^ initial concentration increased tenfold. Even with increasing initial EF-Tu concentration, the IF2·GTP·fMet-tRNA^fMet^ complex, which is required for translation initiation, remains stable.

### 2.4 Converting bell-shaped EF-Tu dependence to a monotonic saturation curve

Using our computational model of the PURE system, we identified a plausible mechanism underlying the bell-shaped EF-Tu dependence. This analysis further suggested a strategy to convert the bell-shaped dependence into a monotonic saturation curve, which we then tested experimentally.

We hypothesized that the bell-shaped profile arises from a shortage of Met-tRNA^fMet^, caused by sequestration of Met-tRNA^fMet^ in the EF-Tu·GTP·Met-tRNA^fMet^ complex. Increasing the initial concentration of tRNA^fMet^ was therefore expected to mitigate this shortage and suppress the bell-shaped dependence. In parallel, raising the concentration of MTF was expected to enhance formylation of Met-tRNA^fMet^ before its interaction with EF-Tu, thereby reducing the accumulation of the non-productive complex EF-Tu·GTP·Met-tRNA^fMet^ complex.

Based on this hypothesis, we examined EF-Tu concentration dependence experimentally under conditions with elevated initial concentrations of both tRNA and MTF. Specifically, the MTF concentration was increased from 0.039 to 0.588 µM, and tRNA from 0.194 to 0.778 µM, while all other components were kept identical to those in Figure 1 (see Methods and Table S1). These concentrations of tRNA and MTF were chosen such that further increases no longer enhanced the amount of peptide synthesis, as determined from titration experiments.

Simulations of the ePURE model using the modified initial concentrations and the fitted parameters described above reproduced the disappearance of the bell-shaped EF-Tu dependence (Figure S5). Consistently, experimental measurements using the PURE system yielded no bell-shaped profile under these modified conditions (Figure 6). These results support the proposed mechanism and suggest that the origin of the bell-shaped EF-Tu dependence identified in ePURE may also operate in the PURE system.

**Figure 6:**
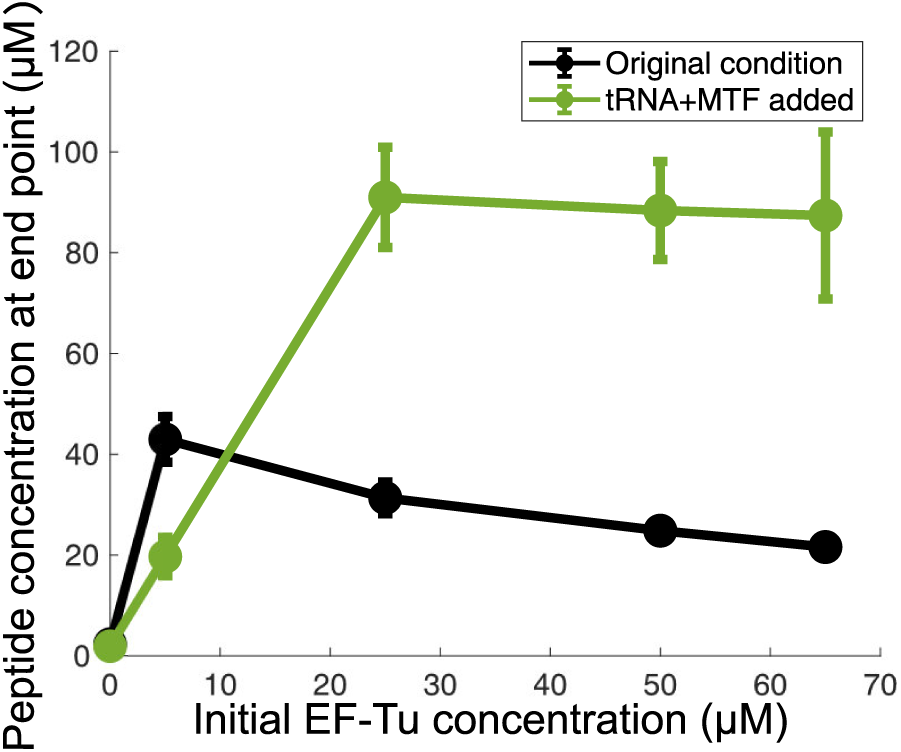
Effect of increasing the tRNA and MTF concentrations on peptide synthesis in the PURE system. Compared with the original experimental results (black; identical to those shown in Figure 1B), the PURE system supplemented with increased concentrations of both tRNA and MTF (green) no longer exhibits the bell-shaped dependence. Error bars represent the standard deviation of four replicate experiments.

## 3 Conclusion

To reconcile discrepancies between experimental observations of the PURE system and the behavior predicted by its mechanistic model, ePURE, we fitted the model’s kinetic parameters using a genetic algorithm. This procedure yielded parameter sets that quantitatively reproduced the experimental time courses. Analysis of these simulations suggested the mechanism underlying the bell-shaped dependence of the peptide production on EF-Tu concentration: excess EF-Tu drives the formation of a non-productive EF-Tu·GTP·Met-tRNA^fMet^ complex, thereby depleting the pool of initiator tRNA available for translation initiation. Guided by this insight, we experimentally altered the concentrations of tRNA^fMet^ and MTF, which eliminated the bell-shaped dependence as predicted.

A central challenge in interpreting complex biochemical systems is that input-output relationships often emerge from distributed and competing interactions rather than from a single rate-limiting step. Even for a well-characterized factor such as EF-Tu, increasing its concentration does not guarantee increased translational output because numerous reactions share substrates, cofactors, and intermediate complexes. Such interdependencies make it difficult to infer causal mechanisms solely from perturbation experiments, particularly in dense reaction networks like those of PURE or living cells.

In this context, a mechanistic framework such as ePURE is essential for disentangling the causal architecture of the system. By explicitly representing all relevant reaction pathways, the model enables systematic evaluation of how perturbations propagate through the network. Our ability to identify tRNA^fMet^ sequestration as a plausible explanation for the bell-shaped response illustrates this strength. Mechanistic modeling thus provides a powerful complement to experimental approaches when interpreting non-intuitive behaviors in cell-free and cellular translation.

Beyond explaining a specific phenomenon, this study highlights a general design principle for engineering cell-free systems. The assumption that elevating the concentration of a single translation factor will monotonically enhance performance does not necessarily hold in complex biochemical networks. Competition for shared molecular resources or the formation of non-productive complexes can generate unexpected non-monotonic outcomes. Identifying components that participate in multiple competing pathways is therefore crucial for rational optimization of translational efficiency.

Despite the successes of the present approach, several technical limitations point toward future model development. Simulations were performed using a short model peptide; extending the framework to longer proteins will require explicit treatment of codon-dependent elongation rates, aa-tRNA demand, and context-dependent ribosomal pausing. Moreover, the current version of ePURE does not incorporate environmental factors such as temperature, magnesium concentration, pH, or ionic strength, all of which strongly influence translation in the PURE system. While incorporating these diverse effects leads to an enormously expanded parameter space, data-driven approaches such as active learning can efficiently guide the parameter exploration and optimization (*27*). Ultimately, incorporating these factors, as well as stochastic fluctuations inherent to systems consisting of molecules with low-copy-number, will be essential for achieving a more complete and predictive description of cell-free translation dynamics, particularly in confined cellular environments.

## 4 Methods

### 4.1 Quantification of protein synthesis by PURE system

The components constituting the PURE system were prepared essentially as described previously (*9*). In brief, proteins were purified via histidine-tag using affinity chromatography, and the ribosome was purified using hydrophobic interaction chromatography and ultracentrifugation. The purchased chemicals with commercial sources and their final concentrations are given in Table S1.

The sequence of plasmid DNA encoding GFP is given in Table S2. Template DNAs for mRNA synthesis were prepared by PCR using KOD Plus Neo (TOYOBO). The PCR products were purified with NucleoSpin Gel and PCR Clean-up kit (Macherey-Nagel) according to the manufacturers’ protocols. mRNA synthesis was carried out using ScriptMAX Thermo T7 Transcription kit (TOYOBO) at 37^◦^C for 2 h. After template DNA digestion with DNase I, mRNA was purified with NucleoSpin RNA Clean-up kit (Macherey-Nagel) according to the manufacturers’ protocols. mRNA concentrations were determined by absorbance measurement at 260 nm.

GFP fluorescence was monitored in real time to quantify protein synthesis in the PURE system reactions. GFP fluorescence measurements were performed using a real-time PCR machine (Mx3005P, Agilent). Reactions were done at 20 µL per well and incubated at 37^◦^C throughout the measurement. 40 nM Alexa647 was incorporated to cancel the well-to-well variation. RNase inhibitor was added to a final concentration of 0.4 U/µL. GFP fluorescence was recorded at 1-min intervals using an excitation wavelength of 492 nm and an emission wavelength of 516 nm. For each reaction condition, four technical replicates were measured, and the mean fluorescence intensity was used for downstream analysis.

### 4.2 Conversion of GFP Fluorescence to Peptide Concentration

To compare the experimental fluorescence trajectories with the ePURE simulations, GFP fluorescence was converted into the concentration of the Met-Gly-Gly tripeptide. Fluorescence intensities were first converted to GFP concentrations using a calibration curve generated from purified GFP standards. Because experiments were performed on two different days with two technical replicates each, inter-day variability was corrected by scaling the fluorescence traces so that the fluorescence values at the EF-Tu concentration of 65 µM matched across days.

The fluorescence signal corresponds to the concentration of folded GFP, whereas the ePURE model does not include folding kinetics. Therefore, we corrected for GFP maturation using a first-order folding model (*28*). The folded GFP concentration *f*^model^[d*p*/d*t*](*t*) was assumed to satisfy the following relationship as a functional defined by the time derivative d*p*/d*t* of the pre-folded concentration *p*(*t*):

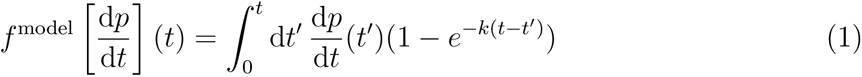

In this equation, the folding rate constant *k* was set to 1/7s^−1^based on reported GFP maturation rates (*29*, *30*). The pre-folded GFP production rate per unit time d*p*/d*t* was estimated using Bayesian inference so that the forward model best matched the experimental folded GFP time series. A non-negative uniform prior was placed on d*p*/d*t*, while the likelihood L[d*p*/d*t*] incorporated Gaussian errors and a smoothness constraint penalizing large second derivatives of *p*(*t*):

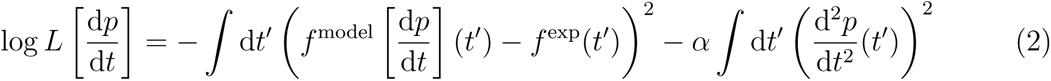

*f*^exp^(t) is the time-series data obtained experimentally. A smoothness constraint and non-negativity were applied to prevent unrealistic fluctuations in *p*(*t*). The smoothness strength *α* was set to 1. Slice sampling generated posterior means from 1,000 MCMC samples (1,000 burn-in) per replicate/EF-Tu condition. By integrating the d*p*/d*t*, the pre-folded GFP concentration *p*(*t*) was obtained.

To relate pre-folded GFP production to synthesis of the 3-amino-acid peptide simulated in ePURE, we assumed that the translation rate is proportional to peptide length. The reconstructed pre-folded GFP concentration was therefore multiplied by 238/3, corresponding to the ratio of the amino acid length of GFP (238 residues) to that of the tripeptide. This conversion was applied independently to all four fluorescence trajectories, and the mean trajectory was used as the experimental reference for kinetic parameter fitting. The experimental data presented in this paper represent the concentrations equivalent to the pre-folded tripeptide converted using the method described in this subsection.

### 4.3 ePURE: Computational model of the translation system

We used a previously developed deterministic ODE model of peptide synthesis based on components of the *E. coli*-derived *in vitro* translation system (PURE system). The model describes the synthesis of the formyl-Met-Gly-Gly (MGG) tripeptide using 241 molecular species in total, including 27 species present initially, and 968 reactions (*15*). All kinetic parameters were obtained from the literature. The kinetic parameters used in this study are listed in Table S4.

In the original ePURE model, 241 molecular species and 968 reactions are included. We removed reactions whose kinetic parameters were set to zero in the original implementation, resulting in a reduced network containing 481 reactions and 205 components. This modification preserved the simulation results exactly while reducing computation time.

### 4.4 Parameter fitting

We fitted the kinetic parameters of the computational model to reproduce the experimental results. A genetic algorithm (GA) was used for parameter fitting (*31*). In the GA, each individual represents a set of kinetic parameters. After introducing random mutations to the parameters, individuals were selected based on the objective function. Details of the GA procedure are described below.

#### 4.4.1 Objective function of GA

The objective function to be minimized is defined as the sum of the squared error between experiments and simulations and a penalty term:

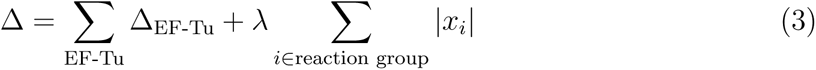

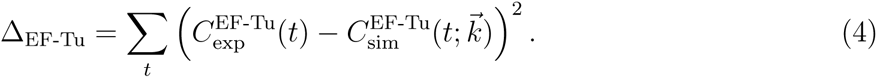

Here, *t* takes every 120 seconds from 120 up to 1,560 seconds and EF-Tu takes 5, 25, 50, and 65 µM, respectively, matching the experimental sampling excluding 0 µM where the simulation result is trivially zero. The vector 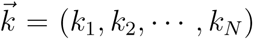 represents the set of *N* kinetic parameter values, and *x_i_* represents the logarithm of the change in *k_i_* as described in the following subsection. 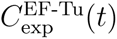 is the experimentally measured peptide concentration under each initial EF-Tu condition, and 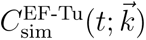 is the corresponding simulated concentration computed using the parameter values 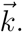

The first term quantifies the agreement between simulation and experiment. The second term penalizes deviations from the original parameter values to maintain biological plausibility. The penalty coefficient was set to *λ* = 1, chosen to be sufficiently large without substantially degrading fitting performance.

#### 4.4.2 Mutation to kinetic parameter values

The ePURE model has the original set of kinetic parameters collected from the literature. Using these values as the initial condition for GA fitting, we introduced mutations according to:

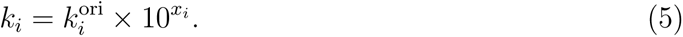

For each reaction i, the kinetic parameter k*_i_*is determined by a deviation x*_i_*from the original parameter 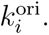 Because kinetic parameters vary widely and often follow a log-normal distribution in biological systems (*32*), deviations were applied on a logarithmic scale.

To ensure biologically reasonable parameters, we imposed the following constraints. First, for reversible reactions, both forward and backward rate constants were changed by the same factor, preserving the equilibrium constant and the associated chemical potentials. Second, reactions sharing the same mechanism and default kinetic parameter values were mutated identically to preserve their equivalence. Examples include the first and second glycine elongation steps in Met-Gly-Gly synthesis, ATP-binding reactions across different processes, and similar phosphate-release reactions (see Table S5 for details). Third, some kinetic parameters in ePURE are assigned very large values to represent sufficiently fast reactions. These parameters were not modified during GA, and we confirmed that allowing them to vary had little effect on simulation outcomes. Finally, Release Factor 1 and 2 (RF1 and RF2) were unified into a single component: all RF2-associated reactions were removed, and the initial RF1 concentration was set to the sum of RF1 and RF2. RF1 and RF2 behave symmetrically in the ePURE model, so distinguishing them adds complexity without providing additional insight. After imposing these constraints, 64 reaction groups remained.

#### 4.4.3 Procedure of GA

The GA was executed as follows:

1. For each individual (parameter set), the translation dynamics were simulated and the objective function was calculated.
2. Among 55 individuals, the 11 with the lowest values of the objective function were selected as parents. Each parent produced 4 offspring via random mutation. Each parameter had a 1% chance of mutation by adding a normally distributed random number to xi in Eq. (5). If no mutation occurred in an offspring, a random mutation was forced at one position to ensure at least one mutated parameter.
3. Parents and offspring were passed to the next generation, and the process was repeated from step 1.

This mutation-selection cycle was repeated for 1,000 generations to obtain fitted parameters. The entire fitting procedure was repeated 100 times using different random seeds to examine the variability of the fitting results (Figure 2A).

## Acknowledgements

This work was supported by RIKEN Junior Research Associate Program (to S. B.), JSPS KAKENHI (22H05403, 24H01118, and 25H01390 to YH; 22K21344, 23K27164, 24H01798 to CF), and GteX Program (JPMJGX23B4 to YH).

## Supporting information

- SupportingFigures.pdf contains the following figures: Figure S1, extended data related to Figure 2A; Figure S2, additional simulation results related to Figure 3B; Figure S3, evidence that IF2·GTP·fMet-tRNA^fMet^ is rate-limiting at high EF-Tu concentrations; Figure S4, accumulation of EF-Tu·GTP·Met-tRNA^fMet^ complexes across all GA-fitted trials; and Figure S5, simulation results corresponding to the additional experiments.
- SupportingTables.xlsx contains the following tables: Table S1, chemicals, their commercial sources, and their final concentrations; Table S2, the sequence of the plasmid DNA pET-G5tag encoding GFP; Table S3, the initial concentrations used in the ePURE simulations, together with the initial concentrations and fluorescence data from two experiments; Table S4, kinetic parameters used in this study; and Table S5, groups of reactions sharing the same mechanism, for which kinetic parameter values were mutated identically to preserve their equivalence.
- The code shows the conversion from fluorescence data to simulation concentrations, the genetic algorithm and its output data, and the output of each figure (https://github.com/ban-shunnosuke/EF-Tu-Bell-Shaped-Translation-Dependency-Analysis).

