## Supporting Figures for "Elucidating the Bell-Shaped Dependence of Protein Translation Activity on EF-Tu Concentration in a Reconstituted Cell-Free System Using a Mechanistic Model"

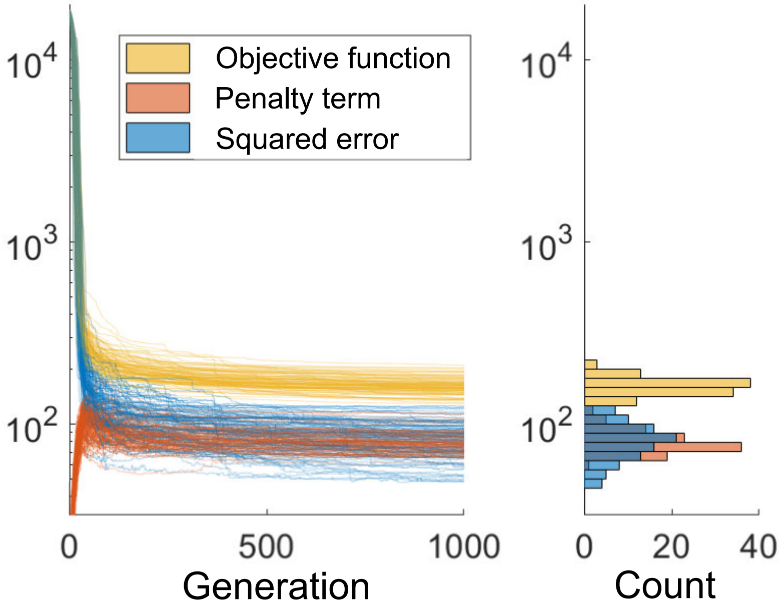


Figure S1: Changes and distribution of squared error, penalty, and the objective function (sum of squared error and penalty) via GA. Change between experimental data and simulations over GA generations (left) and error distribution in the final generation (right).

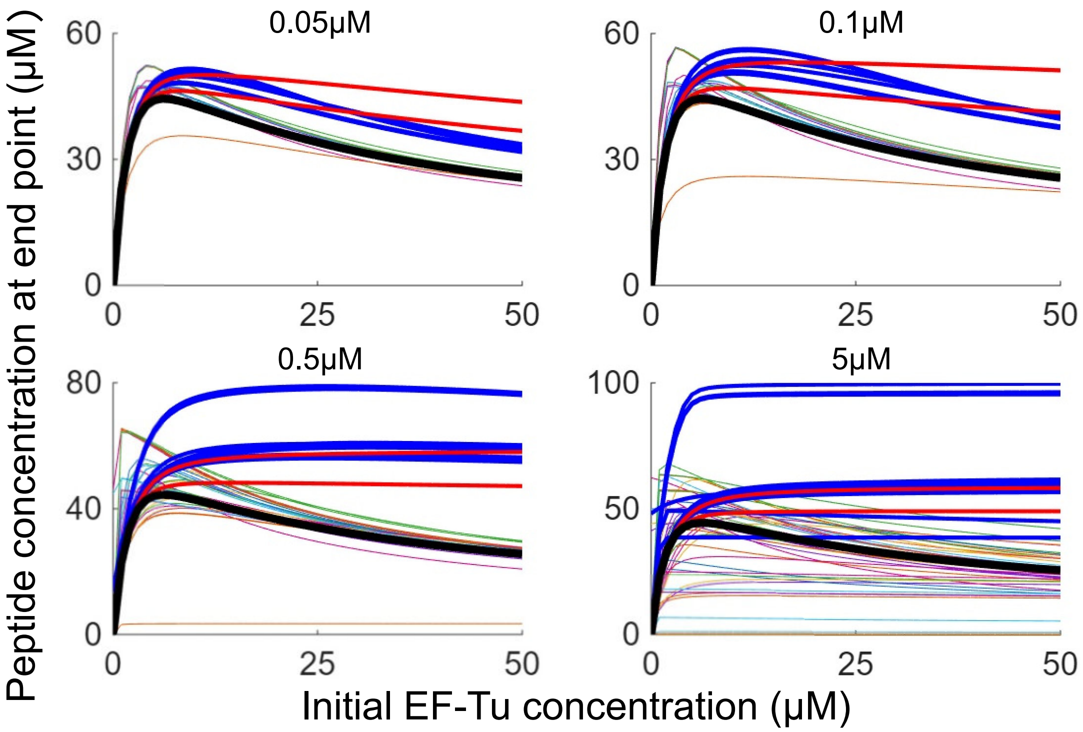


Figure S2: Impact of varying initial concentration increases on initial EF-Tu dependence of Figure 3B. While Figure 3B shows simulations in which the initial concentration of each component was individually increased by 1 µM, this figure displays the results for concentration increases of 0.05 µM, 0.1 µM, 0.5 µM, and 5 µM. Across all tested concentrations, the initial EF-Tu concentration dependence deviates significantly from the baseline (no concentration change, thick black line) when either $tRNA^{fMet}$-containing components (thick blue line) or MTF-containing components (thick red line) are increased.

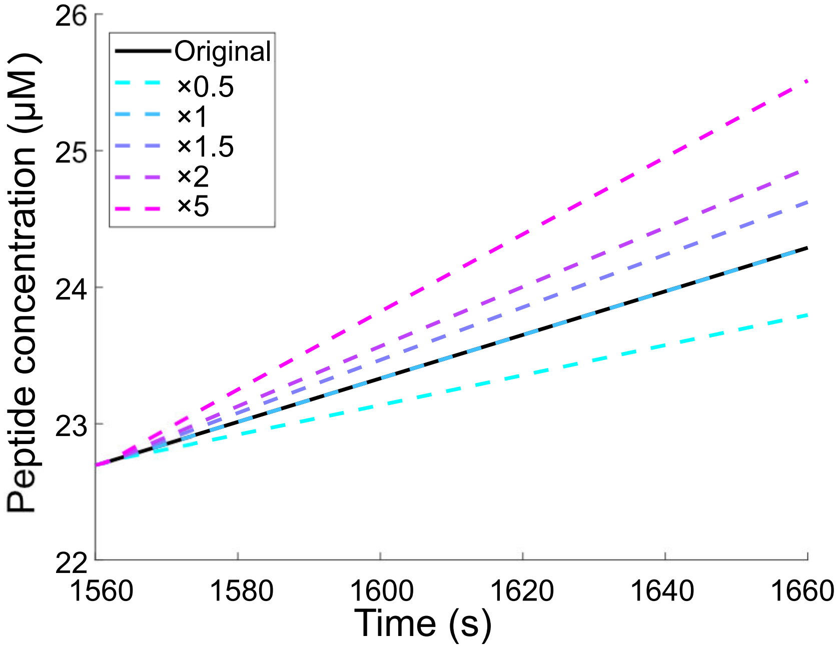


Figure S3. Effect of IF2·GTP·fMet-tRNA$^{\mathrm{fMet}}$ concentration change on peptide production. In a simulation with an initial EF-Tu concentration of 65 μM, for an additional 100$s$ of simulation starting at $t = 1,560s$, the concentration of IF2·GTP·fMet-tRNA$^{\mathrm{fMet}}$ was fixed at specific constant values: 0.5, 1, 1.5, 2, and 5 times its concentration at $t = 1,560s$ (dashed lines, color gradient from cyan to magenta). These perturbed conditions are compared to a original simulation over the same 100-s period (black solid line). The enhanced peptide production resulting from the increased IF2·GTP·fMet-tRNA$^{\mathrm{fMet}}$ concentration demonstrates that this component is rate-limiting.

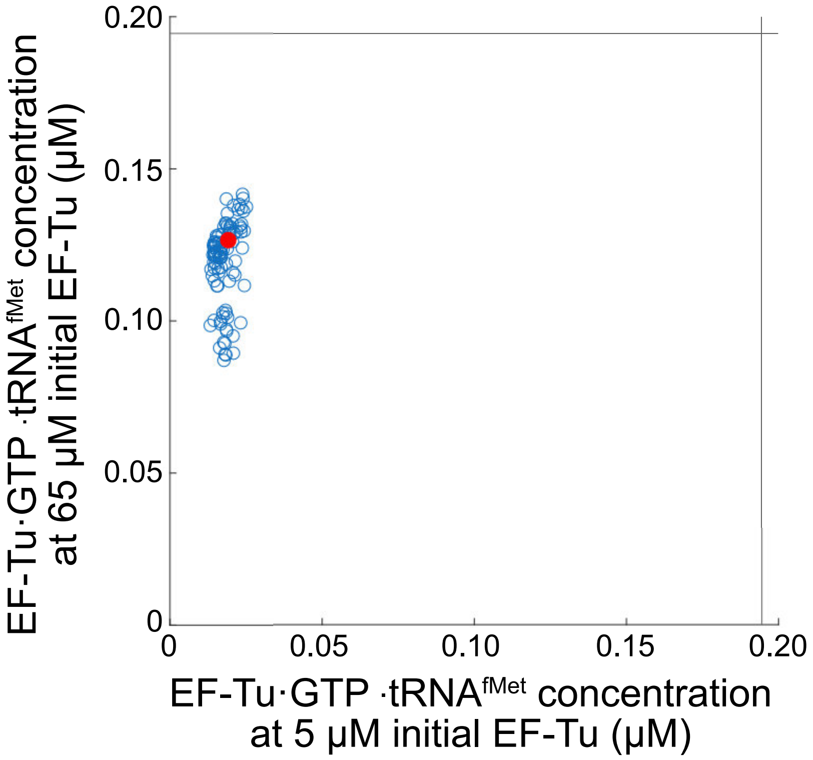


Figure S4: Time-averaged concentration of the EF-Tu·GTP bind Met-tRNA$^{\text{fMet}}$ complex at initial EF-Tu concentrations of 5 μM and 65 μM. Across all GA trials, no accumulation of this complex was observed at 5 μM, while clear accumulation occurred at 65 μM. The black vertical and horizontal lines indicate the total concentration of tRNA$^{\text{fMet}}$-containing complexes, and red dots represent the GA trials used in Figure 4.

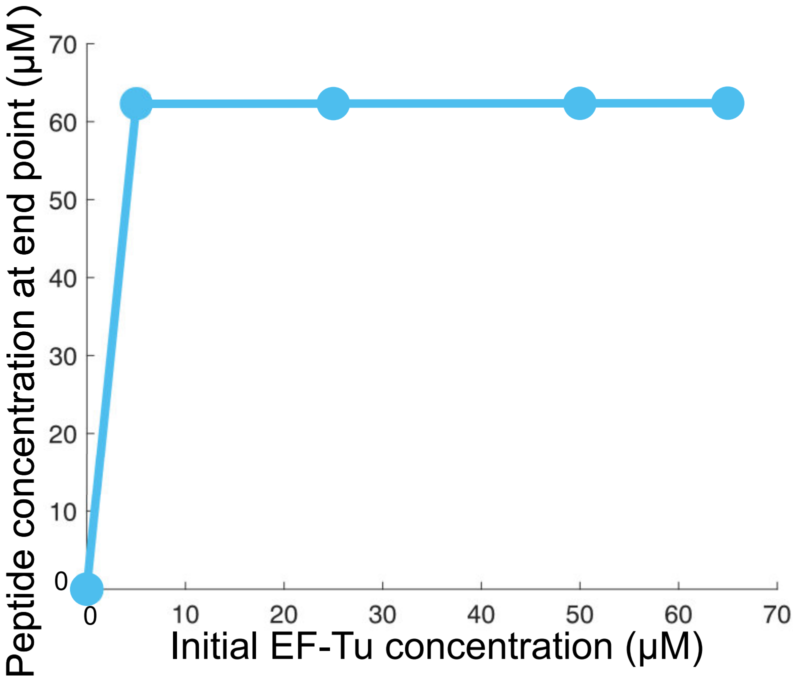


Figure S5: Simulation result which corresponds to the experiments in Figure 6. The initial concentrations of this simulation was set to those identical to the modified initial concentrations of tRNA and MTF used in Figure 6, while the kinetic parameters are identical to those in Figure 2C.
